# p16 expression confers sensitivity to CDK2 inhibitors

**DOI:** 10.1101/2025.02.10.637344

**Authors:** Chance Sine, Lotte Watts, Brianna Fernandez, Jianxin Wang, Erik S. Knudsen, Agnieszka K. Witkiewicz, Sabrina L. Spencer

## Abstract

Blocking the cell cycle is a promising avenue for cancer therapy, with Cyclin-Dependent Kinase 2 (CDK2) emerging as a key target. However, in multiple cell types, CDK4/6 activity compensates for CDK2 inhibition and sustains the proliferative program, enabling CDK2 reactivation. Thus, we hypothesized that sensitivity to CDK2 inhibition is linked to the absence of this CDK4/6-mediated compensatory mechanism. Here we show that Cyclin E1-driven ovarian cancers often co-express the tumor suppressor p16, which inhibits CDK4/6. We show that ovarian cancer cells expressing p16 exhibit heightened sensitivity to CDK2 inhibitors and that depletion of p16 renders them significantly more resistant. Multiplexed immunofluorescence of 225 ovarian patient tumors reveals that at least 18% of tumors express high Cyclin E1 and high p16, a group that we expect to be particularly sensitive to CDK2 inhibition. Thus, p16 may be a useful biomarker for identifying the patients most likely to benefit from CDK2 inhibitors.

## Introduction

Cyclin-Dependent Kinases (CDKs), when complexed with their cognate cyclins, drive cell-cycle entry and progression. Cells initiate the cell cycle through two main pathways: the canonical Cyclin D-CDK4/6-driven pathway^1^ and a less common Cyclin E-CDK2-driven pathway, which bypasses CDK4/6 and is particularly prevalent in cancer contexts^2–4^. Both CDK4/6 and CDK2 phosphorylate the tumor suppressor Rb, leading to the release of the E2F transcription factor, which upregulates genes necessary for S-phase entry^5–7^ (Fig. 1A).

**Fig. 1.**
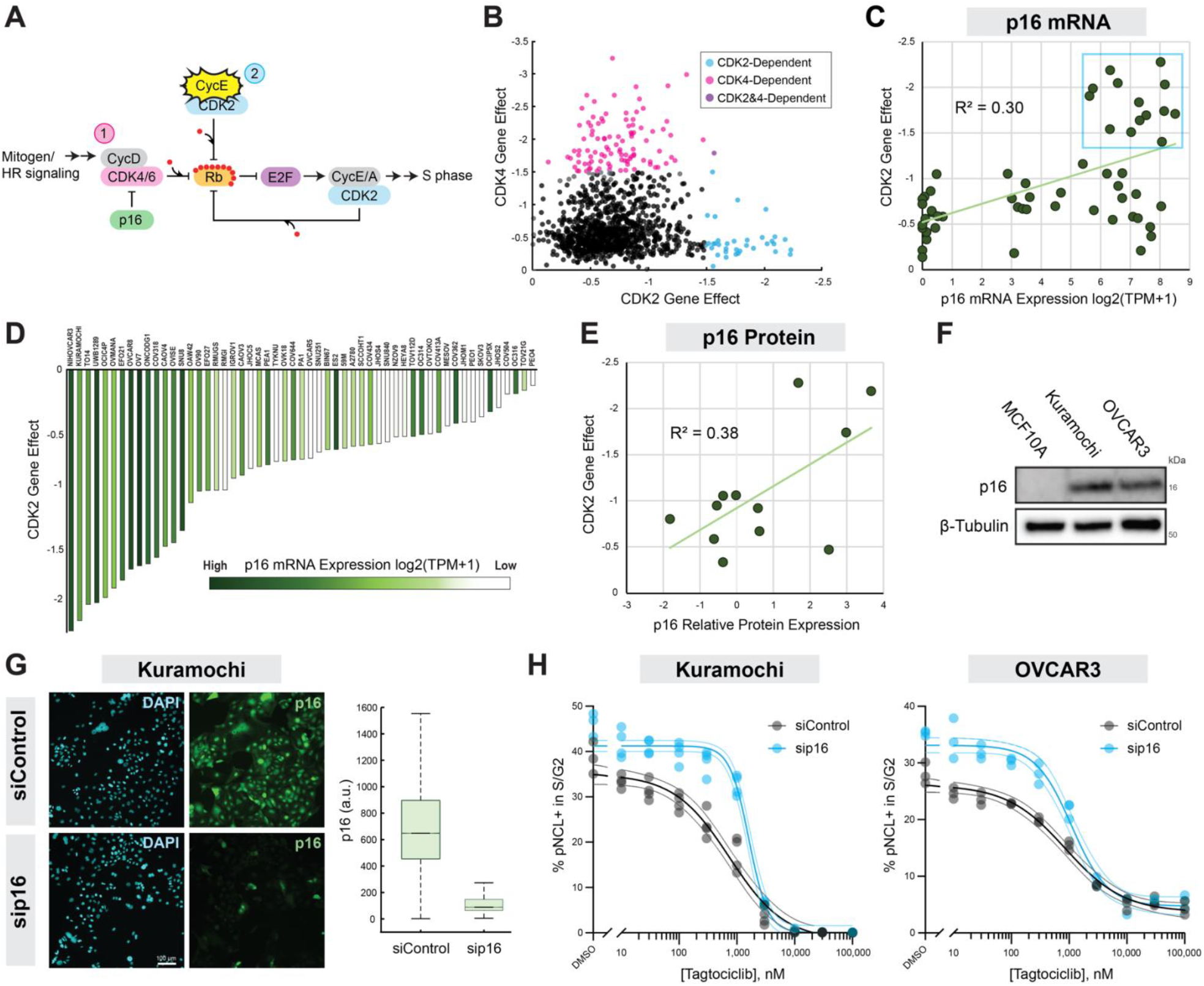
p16 expression is associated with CDK2 dependency in Cyclin E1-amplified ovarian cancers. (**A**) Schematic of normal cell-cycle progression (1) and cell-cycle progression in the Cyclin E-amplified context (2). (**B**) CDK4 dependency vs. CDK2 dependency DepMap data for all cancer cell lines. Dependency scores less than -1.5 are colored as dependent (DepMap Public 23Q4+Score Chronos). (**C**) DepMap CRISPR Gene Effect score (Chronos) for CDK2 vs CDKN2A mRNA expression in ovarian cancers. Linear regression line shown in light green along with R^2^ value. Blue square highlights that all CDK2-dependent lines have high p16 mRNA. (**D**) Waterfall plot ordered on CDK2 Gene Effect score (Chronos) and colored by p16 mRNA expression in ovarian cancer cell lines. Each bar is labeled with the cell line name. (**E**) DepMap CRISPR Gene Effect score (Chronos) for CDK2 vs. p16 relative protein expression in ovarian cancers. Linear regression line shown in light green along with R^2^ value. (**F**) Western blot assessing p16 protein expression in MCF10A, Kuramochi, and OVCAR3. (**G**) p16 immunofluorescence in Kuramochi with and without p16 siRNA knockdown for 72h. (**H**) Kuramochi and OVCAR3 were treated with the indicated dose curves of Tagtociclib for 8 h after which cells were fixed and processed for immunofluorescence. Dose-response curves show the percent of phospho-NCL+ cells in S/G2 (determined by DNA content). Three technical replicates per condition, each individual replicate shown. Four-parameter inhibitory Hill equation was employed in GraphPad Prism to fit the curves. Outside lines indicate 95% confidence interval; the IC50 in Kuramochi is 742nM (siControl) vs. 1612nM (sip16) in and in OVCAR3 is 861nM (siControl) vs. 1102nM (sip16).

Dysregulation of proliferation driven by CDK activity is a hallmark of cancer, making CDKs prime targets for therapeutic intervention aimed at halting aberrant proliferation. Among the CDK family members, CDK2 stands out for its pivotal role in regulating the Restriction Point, the moment of cell-cycle commitment^8–10^. In recent years, there has been increasing attention on developing CDK2 selective inhibitors as potential cancer therapies. A motivation for the development of CDK2 inhibitors were patients with hormone receptor (HR)-positive/human epidermal growth factor receptor 2 (HER2)-negative breast cancers that had developed resistance to CDK4/6 inhibitors via upregulation of Cyclin E-CDK2 activity^11,12^. Now, numerous CDK2 inhibitors are also advancing through clinical trials as a primary treatment option for a variety of solid tumor types, including those with amplified Cyclin E1 (gene name *CCNE1*)^13–18^.

CCNE1-amplified ovarian cancers, representing 15-20% of all ovarian cancers^19–21^, are a population with unmet clinical need due to their resistance to chemotherapy and poor prognosis^22,23^. Further, CCNE1 amplification is mutually exclusive with BRCA1/2 and other DNA damage repair-related mutations^24^, leading to poor outcomes with therapies that target these DNA damage repair deficiencies, such as PARP inhibitors or platinum-based therapies^25–27^.

Recent work from our lab has shown that when CDK2 is inhibited, many cell types are able to compensate for the loss of CDK2 activity, utilizing a CDK4/6-dependent mechanism that enables cells to continue cycling^28^. More specifically, under CDK2 inhibition (CDK2i), CDK4/6 can maintain Rb phosphorylation and liberation E2F to drive production of Cyclins E and A, leading to a rebound in CDK2 activity^28^. However, ovarian cancers with amplified CCNE1 are known to be particularly sensitive to CDK2 inhibitors^25,29–31^. The reason for this sensitivity is thought to be that inhibition of the ‘driver’ of these ovarian cancers, Cyclin E1, by CDK2 inhibitors blocks proliferation. However, one would expect that these cells should be primed for incomplete CDK2 inhibition enabling CDK2 reactivation due to their already high baseline CDK2 activity, and potential access to the same CDK4/6-dependent adaptive mechanism that we observed in other cell types^28^. We therefore hypothesized that something must be blocking CDK4/6 from compensating for the loss of CDK2 activity in CCNE1-amplified ovarian cancers.

Interestingly, CCNE1-amplified ovarian cancers often express high levels of the tumor suppressor p16^INK4A^ (p16, gene name *CDKN2A*), which inhibits CDK4/6 activity^32^. Consistent with p16’s role as a cell-cycle inhibitor, its expression is often used as a marker of senescence and is rarely expressed in normal proliferative tissues^33^. This paradoxical expression of p16, a cell-cycle inhibitor, in fast-cycling ovarian cancer cells is not understood.

In this study, we investigated the ability of p16 to modulate the sensitivity of CCNE1-amplified ovarian cancer cells to CDK2 inhibitors. Our findings reveal that cells expressing p16 exhibit heightened sensitivity to CDK2 inhibitors due to p16’s inhibition of CDK4/6, which prevents cells from compensating for the loss of CDK2 activity. This mechanistic insight suggests a promising therapeutic strategy that leverages p16 status to tailor CDK2-targeted treatments to the subset of patients that are most likely to respond to this new class of drugs.

## Results

### DepMap analysis reveals that cells expressing p16 have an increased dependency on CDK2

Consistent with the concept that there are two cell-cycle entry pathways (Fig. 1A), genome-wide CRISPR knockout viability screens show that cells can be reliant on CDK2 or CDK4/6 for proliferation, but not both ^34,35^ (Fig. 1B). However, the factors that determine sensitivity or resistance to perturbations of each pathway remain unclear. To address this, we examined predictors of CDK2 dependence using DepMap gene-dependency data and identified CDKN2A (p16) expression as a significant factor (Fig. S1A). Notably, while heightened CCNE1 mRNA expression or gene amplification is known to be associated with CDK2 dependence, CCNE1 expression is less predictive of CDK2 dependency than p16 expression^34,35^.

The plurality of CDK2-dependent cell lines in DepMap are derived from ovarian cancers (Fig. S1B), which prompted us to focus on this cancer type for our study. Our further analysis revealed that all CCNE1 -amplified ovarian cancers in the DepMap database also exhibit high p16 mRNA expression (Fig. S1C).

Additionally, ovarian cancer lines with high CDK2 dependence consistently show elevated p16 mRNA and protein levels (Fig. 1C blue box, 1D-E). Interestingly, we did not find the same trend with any of the other CDK inhibitor proteins (Fig. S1D). Based on these data from DepMap and informed by our recent work on CDK2 inhibitors^28^, we hypothesized that the sensitivity of these cancers to CDK2 inhibitors is not solely attributable to elevated Cyclin E1 expression, the current model in the field. Instead, p16 expression may play a critical role by inhibiting CDK4/6, thereby preventing these cells from compensating for CDK2 inhibition via use of CDK4/6.

### Cells expressing high p16 are sensitive to CDK2 inhibition

To test the hypothesis that p16 expression modulates sensitivity to CDK2 inhibitors, we focused on two Cyclin E1-driven, high-grade serous ovarian cancer cell lines, OVCAR3 and Kuramochi. Western blot analysis confirmed that both OVCAR3 and Kuramochi cell lines express p16, while non-transformed MCF10A mammary epithelial cells, which have a p16 deletion, do not (Fig. 1F).

We next performed p16 knockdown experiments in OVCAR3 and Kuramochi cells to assess the impact of reduced p16 levels on CDK2 inhibitor sensitivity. Following successful p16 knockdown (Fig. 1G and Fig. S1E), cells were treated with the CDK2 inhibitor Tagtociclib (formerly known as PF-07104091 or PF4091), which is currently in phase 1 and 2 clinical trials, including trials in ovarian cancer^13^. We assessed the dose response to Tagtociclib using single-cell immunofluorescence to measure the CDK2 substrate Nucleolin^36^ (pNCL, phosphorylated at T84), in both p16-expressing and p16-depleted cells. Our analysis revealed a rightward shift in the IC50 for Tagtociclib in p16-depleted cells compared to p16-expressing cells. This result suggests that p16 depletion renders cells more resistant to CDK2 inhibition (Fig. 1H). Furthermore, these data indicate that p16 expression is not merely associated with CDK2 dependence but plays a causal role in modulating the cellular response to CDK2 inhibitors.

### Live-cell imaging reveals that p16 depletion allows cells to rebound out of CDK2i

We previously utilized single-cell time-lapse imaging with a DNA helicase B (DHB)-based CDK2 activity sensor^9^ (Fig. 2A) to assess the cellular response to Tagtociclib, Palbociclib (a CDK4/6 inhibitor), and their combination. As previously reported, while treatment of MCF10A cells with Tagtociclib leads to an immediate reduction of CDK2 kinase activity, these cells exhibit a rapid rebound in CDK2 activity driven by a CDK2/4/6-Rb-E2F-dependent mechanism that circumvents CDK2 inhibition and enables cell-cycle completion (Fig. 2B)^28^. By contrast, combining Tagtociclib with Palbociclib, inhibiting both CDK2 and CDK4/6, leads to a sustained suppression of CDK2 substrate phosphorylation and results in cell-cycle exit, as seen previously^28^ (Fig. 2B, Fig. S2A).

**Fig. 2.**
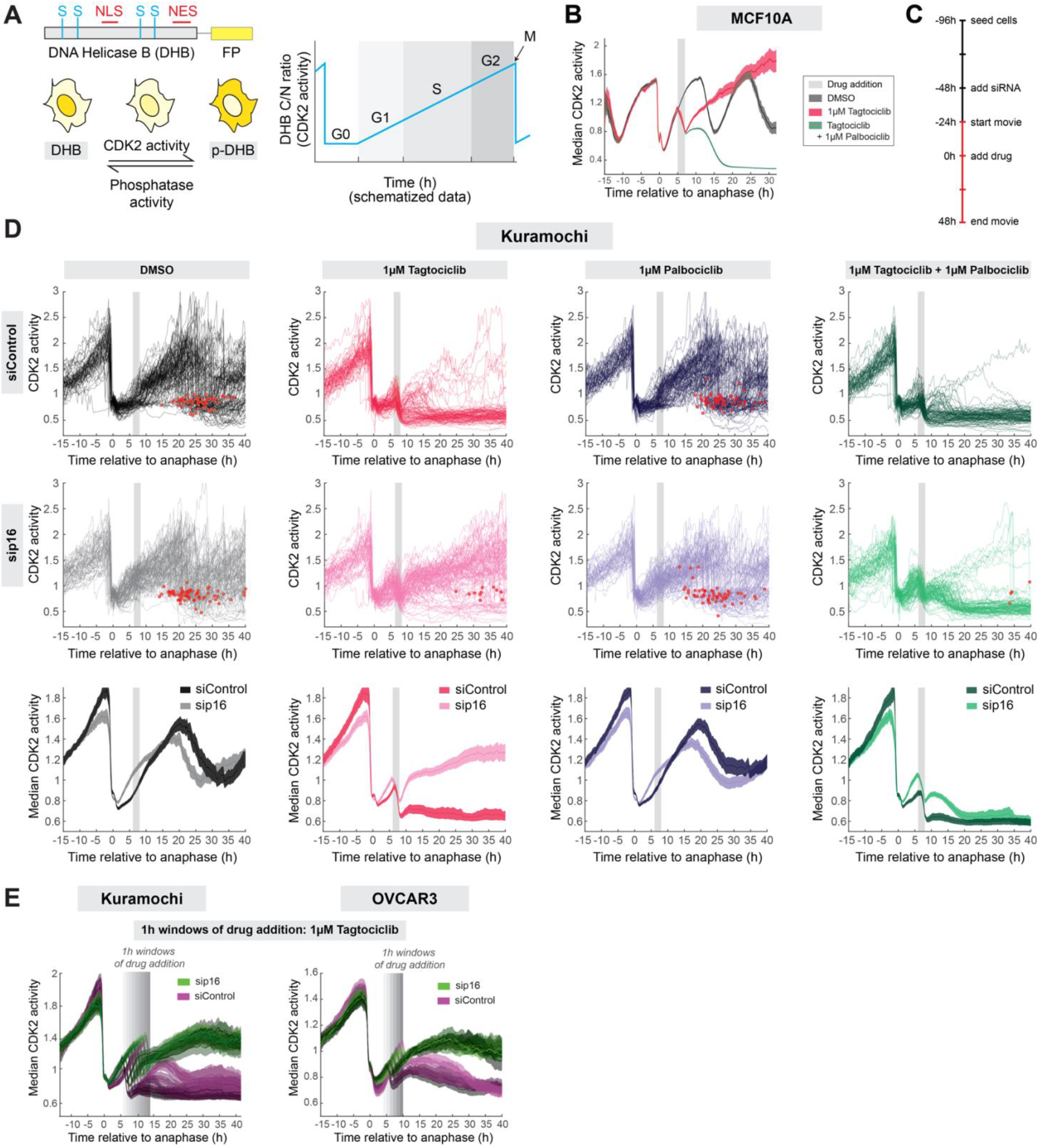
p16 expression modulates the CDK2 activity rebound upon CDK2 inhibition. (**A**) Left: Schematic of the DHB-based CDK2 activity sensor. The sensor localizes to the nucleus when unphosphorylated; phosphorylation by CDK2 leads to translocation of the sensor to the cytoplasm. NLS, nuclear localization signal; NES, nuclear export signal; S, CDK consensus phosphorylation sites on serine. Right: schematized data showing the dynamics of CDK2 activity across G0, G1, S, G2, and M phases of the cell cycle. (**B**) CDK2 activity in MCF10A 2A7T (Cyclin D-mCitrine) cells treated with 1mM Palbociclib, 1mM Tagtociclib, or the combination. Unless stated otherwise, all time-lapse plots are median ± 95% confidence interval as shaded bands where non-overlapping shading indicates a statistically significant difference with a p value < 0.05 (**C**) Timeline for live cell movie with p16 knock down, red indicates the imaging period. (**D**) Top two rows: Single-cell traces of CDK2 activity in Kuramochi cells treated with the indicated drugs. Traces are computationally aligned to the point of anaphase and CDK2^inc^ cells (see Methods) are selected for plotting if they received drug 6-8 h after anaphase (gray bar). Red dots indicate mitosis. Bottom row: overlaid 95% confidence interval for the respective plots above. (**E**) 95% confidence interval for OVCAR3 and Kuramochi cells treated with 1mM Tagtociclib at multiple 1h windows of drug addition (gray bars).

To test whether p16 expression modulates this CDK2 activity rebound in response to CDK2i in Cyclin E1-driven ovarian cancer models, we expressed the CDK2 activity sensor in Kuramochi and OVCAR3 cells. p16-expressing and p16-depleted cells were imaged for 24 hours to establish baseline CDK2 activity before Tagtociclib, Palbociclib, or the combination were added (Fig. 2C). CDK2 activity traces were aligned to the time of anaphase and cells were plotted if they received the drug during G1 phase, 6-8 hours after anaphase (Fig. 2D, Kuramochi; Fig. S2C, OVCAR3). Notably, in the DMSO control condition, cells with p16 knockdown (grey traces) enter the cell cycle more quickly and exhibit a more rapid increase in CDK2 activity that lasts for the majority of the cell cycle. Furthermore, cells with p16 depletion do not elevate CDK2 activity to the same level as control cells prior to mitosis (Fig. 2D, column 1). Despite these alterations in cell-cycle dynamics, p16-depleted cells maintain a similar proportion of cells successfully undergoing mitosis compared to control (Fig. S2B).

As anticipated, due to their known dependence on CDK2, Kuramochi and OVCAR3 cells exhibit sustained suppression of CDK2 activity following treatment with Tagtociclib (Fig. 2D, column 2). Strikingly, depletion of p16 dramatically alters this response, converting the sustained suppression of CDK2 activity into a drop-rebound phenotype like that observed in MCF10A cells (Fig. 2D, column 2). We also observed this striking difference in response to CDK2i across the entire cell cycle, not just in G1 (Fig. 2E). Importantly, p16-depleted cells treated with Tagtociclib are still able to undergo mitosis, whereas control cells treated with Tagtociclib rarely do (Fig. S2B). These data demonstrate that p16 expression modulates the response to CDK2 inhibition in Cyclin E1-driven ovarian cancer cell lines.

Consistent with p16’s role in inhibiting CDK4/6, the addition of Palbociclib does not significantly affect cell-cycle progression in p16-expressing cells (Fig. 2D column 3). Moreover, p16 depletion does not convert cells to being sensitive to CDK4/6i on the timescales tested here, likely due to their cell cycle being driven exclusivity by CDK2 activity prior to acute p16 depletion. The combination of Tagtociclib and Palbociclib more effectively suppresses CDK2 activity in p16-depleted cells, highlighting the necessity of inhibiting total Rb-kinase activity (CDK4/6 and CDK2 activity) to impede cell-cycle progression (Fig. 2D column 4). The observation that cells with p16 depletion rebound in response to Tagtociclib, but not in combination with Palbociclib, suggests that their ability to rebound is due to compensatory CDK4/6 activity rather than an unknown role of p16 in regulating this response.

### p16 levels also modulate phosphorylation of additional CDK2 substrates as well as long-term drug responses

To determine whether our observations are specific to the DHB-based CDK2 sensor or if they extend to other CDK2 substrates, we next investigated the rebound phenotype by measuring the CDK2 substrate, Nucleolin. Consistent with our live-cell imaging results, we observed that control cells exhibit a sustained reduction in pNCL upon treatment with Tagtociclib, while cells with p16 depletion show an initial reduction in pNCL followed by a rebound starting 6 hours after drug addition (Fig. 3A). Unexpectedly, we observed that p16 knockdown leads to an increase in baseline pNCL in DMSO-treated cells, suggesting that p16 is indirectly holding back CDK2 activity via inhibition of CDK4/6 activity, as also seen in Fig. 2D, column 1.

**Fig. 3.**
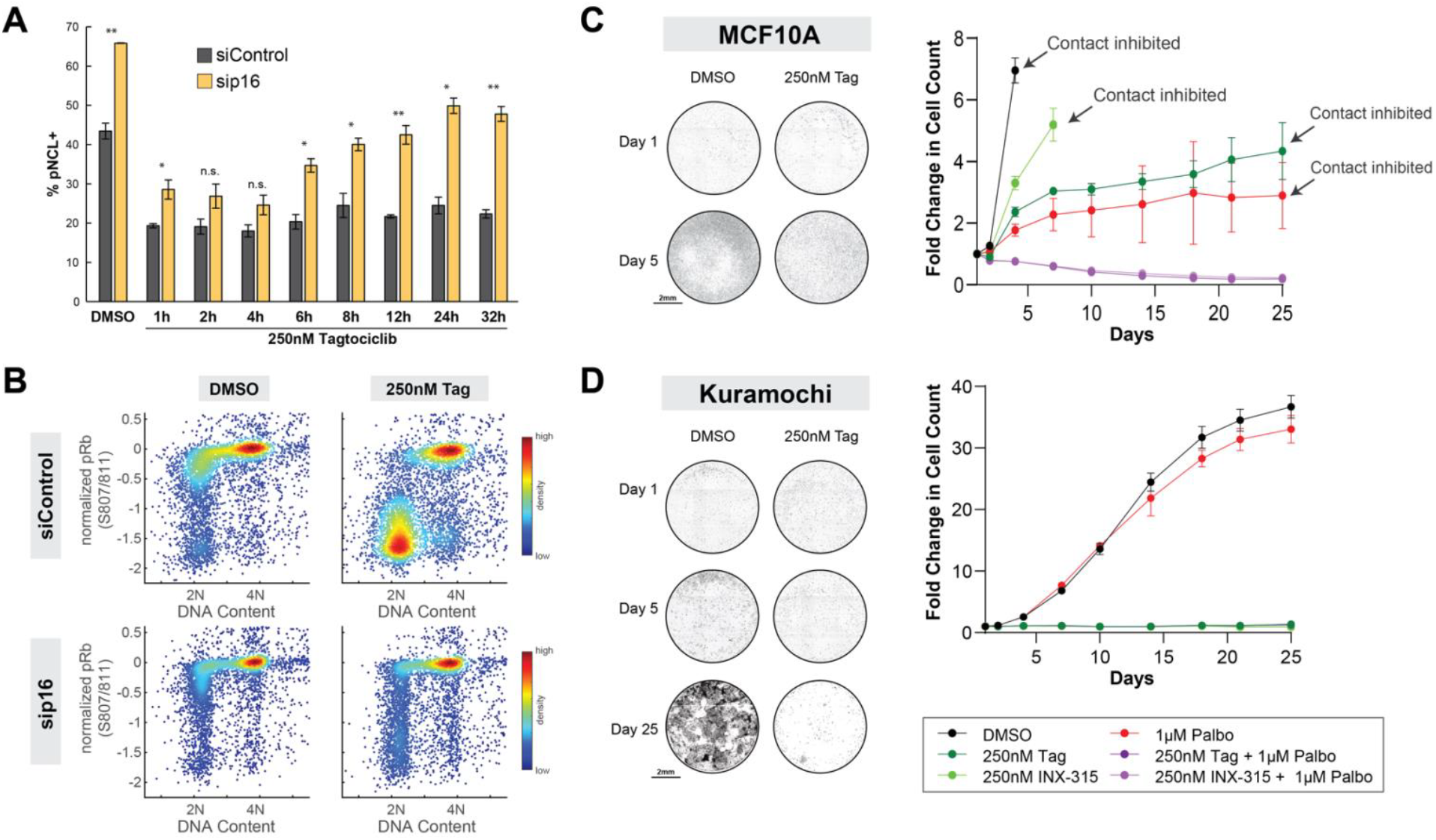
p16 expression modulates CDK2 substrate phosphorylation and cell outgrowth upon CDK2 inhibition. (**A**) Bar charts quantifying the fraction of pNCL-positive Kuramochi cells following treatment with 250nM Tagtociclib for the indicated times. Mean of at least 3 technical replicates plotted ± s.d. (**B**) Density scatter plots portraying the effect of the indicated compound on phospho-Rb at S807/811 in OVCAR3 cells 8h after drug treatment. phospho-Rb was normalized to total Rb in each cell. Note that the top mode of phospho-Rb represents hyperphosphorylated Rb. (**C-D**) Left: Representative images from Day 0 and 5 of MCF10A and Day 0, 5, and 30 of Kuramochi cells treated with the indicated drugs. Right: Quantification of imaging data. Mean of at least 3 technical replicates plotted ± s.d. Note: increased cell size upon treatment with Tagtociclib or Palbociclib results in differing cell counts in which conditions reach contact inhibition.

Similarly, p16 knockdown in DMSO-treated cells caused increased Rb phosphorylation, particularly in cells with 2N DNA content, consistent with CDK4/6’s role in Rb phosphorylation during G1 phase (Fig. 3B column 1). Following an 8-hour treatment with Tagtociclib, p16-depleted cells maintain higher Rb phosphorylation compared to the siControl, aligning with live-cell imaging data showing continued cell cycling under CDK2 inhibition in p16-depleted cells (Fig. 3B column 2).

We next investigated whether p16 expression continues to affect sensitivity to CDK2 inhibitors in the long term. Consistent with previous findings, MCF10A cells lacking p16 proliferate under CDK2 inhibition and require the combination of CDK2/4/6 inhibition to halt growth (Fig. 3C). In contrast, OVCAR3 and Kuramochi cells, which express p16, do not outgrow over 25 days, consistent with their short-term sensitivity to CDK2 inhibitors alone (Fig. 3D, Fig. S3A). Additionally, these p16-expressing cell lines do not respond to CDK4/6 inhibitors, further evidence that they progress through the cell cycle independently of CDK4/6. Overall, assuming the response of MCF10A cells to CDK2i is representative normal cells in the body, these results suggest the existence of a therapeutic window for the use of CDK2 inhibitors in Cyclin E1-driven ovarian cancer.

### Identifying Cyclin E1- and Cyclin D1-driven tumor populations

Following our findings in cell lines, we wanted to determine whether p16 expression also occurs in ovarian patient tumors and how often it is co-expressed with Cyclin E1 and Cyclin D1. While RNA-sequencing data are now ubiquitous in the cancer literature, the cell cycle is one area where the proteome is much more informative than the transcriptome, given that many key regulatory mechanisms are post-translational.

We obtained formalin-fixed paraffin-embedded tumor sections from 225 ovarian cancer patients, which we stained with antibodies against p16, Cyclin E1, Cyclin D1, CDK2, Rb, Geminin, and cytokeratin AE1/AE3. We used Geminin to mark cycling cells, since it is absent in G0 and G1, begins rising at the G1/S transition and increases steadily throughout interphase before being degraded in anaphase^37,38^. AE1/AE3 is used to distinguish tumor from non-tumor cells. Two example tumors are shown in Fig. 4A-C: tumor #19 with high p16, high Cyclin E1, and low Cyclin D1, and tumor # 208 with low p16, low Cyclin D1, and high Cyclin D1. In addition, CDK2 protein levels were elevated in tumor #19 compared to tumor #208 (Fig. 4A-C). In both samples, many tumor cells are Geminin-positive and thus are cycling, even in the presence of high p16 (Fig. 4A-C).

**Fig. 4.**
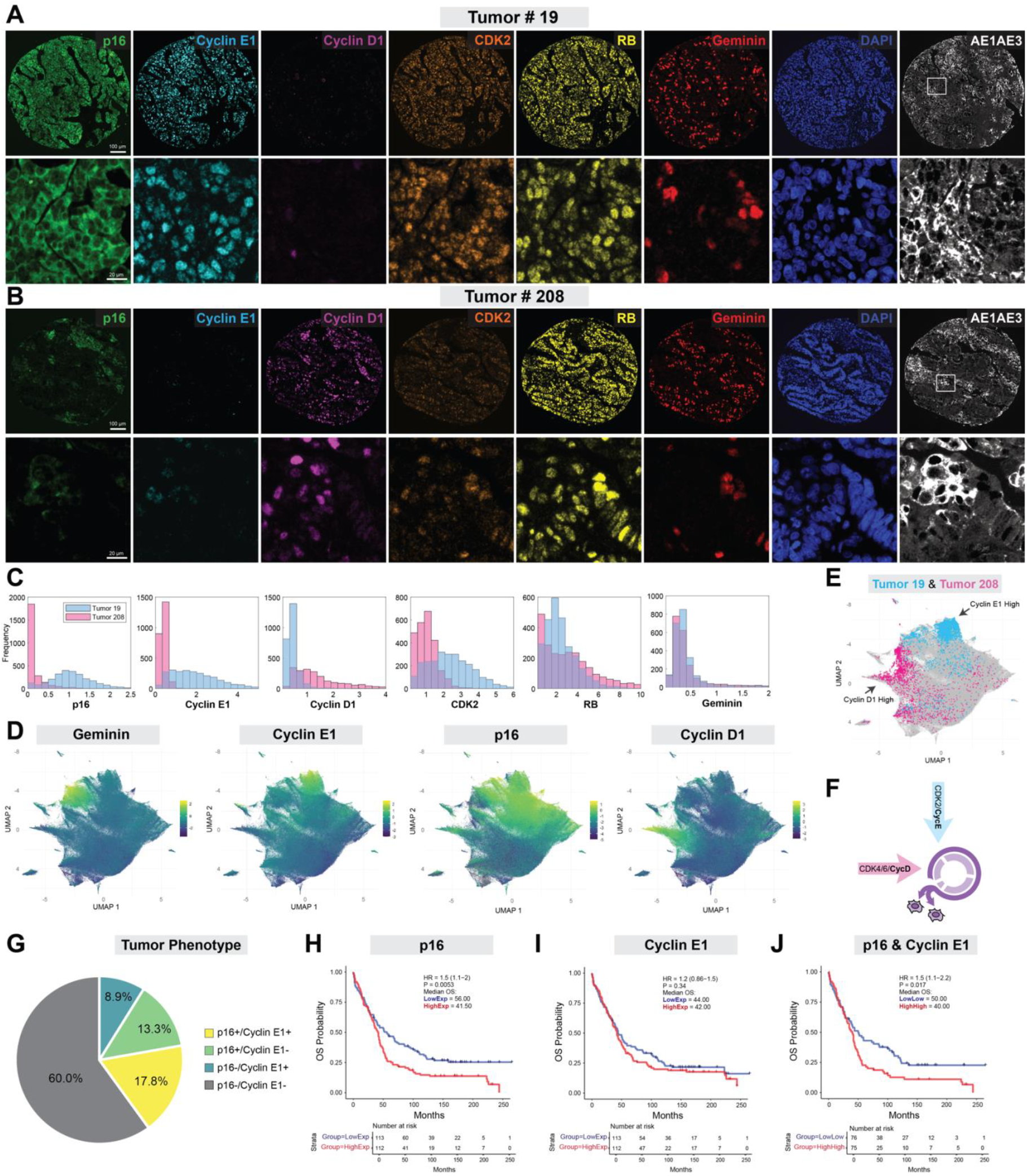
The ovarian tumor cell cycle is driven by either Cyclin E or Cyclin D. (**A-B**) Ovarian tumor cores #19 (A) and #208 (B) stained using a multispectral immunofluorescent (mIF) staining process for p16, Cyclin E1, Cyclin D1, CDK2, total Rb, Geminin, cytokeratin AE1AE3 to mark tumor cells, and DAPI to mark nuclei. Note that the vast majority of cells in the zoomed-in image are tumor cells. (**C**) Single-cell distribution of the intensity of each respective marker in tumor #19 vs. tumor #208. (**D**) Cells from all 225 tumors were pooled to generate a single-cell UMAP from 216 parameters from the 7 markers, colored by the scaled intensity of indicated marker. (**E**) UMAP of cells from tumor core #19 (blue) and #208 (pink), each residing in different regions of the UMAP. (**F**) Schematic showing a Cyclin D-driven and a Cyclin E-driven pathway into the ovarian cancer cell cycle. (**G**) Phenotyping of ovarian patient tumors based on p16 and Cyclin E1. Intensity cutoffs were made using Phenoptr and the most abundant cell phenotype is used to determine tumor phenotype. (**H**) Kaplan-Meier plot by p16 dichotomy. (**I**) Kaplan-Meier plot by Cyclin E1 dichotomy. (**J**) Kaplan-Meier plot by p16-high/CyclinE1-high dichotomy vs. p16-low/Cyclin E1-low dichotomy.

In agreement with cell-line data from DepMap, p16 expression and Cyclin E1 expression are also correlated in our tumor samples (Fig. S4C, D-E). Furthermore, increasing p16 protein expression in the 225 tumors is correlated with increasing Geminin staining and hence cell proliferation, likely enabled by increased Cyclin E1 and CDK2 protein (Fig. S4A-B). We next examined the individual tumor cells by pooling all single cells across all tumors and plotting UMAPs of marker intensity and localization (e.g. nuclear, cytoplasmic, membrane) as well as cell size. This allowed us to visualize where each tumor cell is positioned in high-dimensional cell-cycle space (Fig. 4D). We noted that the Cyclin E1-high cells are positioned together at the top of the UMAP, and that essentially all Cyclin E1-high cells are also p16-high. However, not all p16-high cells are Cyclin E1-high, perhaps because Cyclin E1 is degraded each cell cycle at the G1/S transition^38–40^. This also likely explains why the Cyclin E1-high cells are rarely Geminin-high: Cyclin E1 levels fall at the G1/S transition, the moment when Geminin levels begin to rise^38^. We also noted that Cyclin D1-high cells are positioned together in a peninsula on the left side of the UMAP, and that these Cyclin D1-high cells have reduced p16 and low Cyclin E1 protein expression. Cyclin D1-high cells are also rarely Geminin-high because Cyclin D1 is degraded in S phase when Geminin levels begin to rise^38,41,42^.

In a normal cell cycle, Cyclin D1 and Cyclin E1 levels are both high early in the cell cycle and low in S phase^40–43^. Thus, the fact that the patient tumor cells quantified here generally fall into Cyclin E1-high or Cyclin D1-high groups (e.g., tumor #19 vs tumor #208, Fig. 4E) suggests that these tumor cells utilize distinct paths into the cell-cycle (Fig. 4F). Zooming out to the individual tumor level, our results suggest that different patient tumors may have different pathway utilizations and hence vulnerabilities. Indeed, after determining intensity cutoffs using Phenotypr, we classified each tumor as p16 high/low and Cyclin E1 high/low. We found that 17.8% of tumors express high p16 and high Cyclin E1 (Fig. 4G), although this number is likely an underestimate since Cyclin E1 is degraded in S and G2 phases^38^. Importantly, this p16-high/Cyclin E1-high population is one we propose might be especially vulnerable to CDK2i, since p16 will prevent the CDK4/6-dependent rebound in CDK2 activity.

### Ovarian cancer patients with p16-expressing tumors have poor overall survival

Further analysis of the 225 ovarian patient datasets revealed that patients with p16-expressing tumors experience significantly poorer overall survival (OS) compared to those with low p16 expression (Fig. 4H). When evaluating the impact of Cyclin E1 expression, another established marker of poor OS^44,45^, we found that Cyclin E1-high tumors are not significantly associated with poor OS, and that p16 expression is a more robust predictor of survival outcomes (Fig. 4I). Tumors that are both p16-high and Cyclin E1-high have a somewhat worse prognosis compared to gating on p16-high alone, although the p-value is less significant (Fig. 4J).

These observations highlight a critical population of patients with an unmet clinical need, as high p16 expression is associated with a more aggressive disease course. Given that most of these patients received standard-of-care chemotherapy, our findings suggest that tumors with high p16 expression would require different therapeutic strategies for this patient subgroup, such as CDK2 inhibition.

## Discussion

Our study elucidates the critical role of p16 in modulating sensitivity to CDK2 inhibitors in Cyclin E1-driven ovarian cancers. We demonstrate that high p16 expression, rather than CCNE1-amplification alone, is a key determinant of CDK2 inhibitor sensitivity in these cancers. This finding underscores a nuanced understanding of how p16 impacts cell-cycle regulation and therapeutic response in cancer cells.

p16 is a well-known CDK4/6 inhibitor protein that typically exhibits low expression in normal proliferating tissues^46^, where its role as a marker of cellular senescence is well-documented^33^. Our results show that p16 expression is elevated in CCNE1-amplified ovarian cancers, and that these cells are cycling rather than senescent. We further show that p16 expression in Cyclin E1-high ovarian cancer cell lines enhances sensitivity to CDK2 inhibitors. This sensitivity is due to p16’s inhibition of CDK4/6, which prevents cells from using CDK4/6 to compensate for the loss of CDK2 activity. This interplay of CDK4/6 and CDK2 activity highlights the necessity of inhibiting the totality of Rb-kinase activity to effectively suppress cell-cycle progression^28^.

The finding that p16 sensitizes cancer cells to CDK2 inhibition raises the question of whether loss of p16 will become a mode of resistance to CDK2 inhibition. Indeed, a recent study found that Kuramochi cells, which are resistant to the PARP inhibitor, Olaparib, lose p16 as they become resistant^47^. However, this result also raises the question of why these cancer cells express p16 in the first place. One possibility is that p16 expression is needed to hold back the unscheduled cell-cycle entry and replication stress that is associated with high levels of Cyclin E1^48,49^, a hypothesis that could be tested in future work. Another open question is whether CCNE1 amplification or overexpression is required for p16 expression to render cancer cells sensitive to CDK2 inhibitors. It seems likely that something driving cell-cycle entry via the CDK2 path is required in addition to p16 expression, or else a CDK2 inhibitor may not have any effect, but this requires rigorous examination.

Our findings may not be limited to ovarian cancer. The mechanisms we identified here may extend to other cancer types characterized by high Cyclin E1 and high p16 expression. For example, some gastric cancers have shown similar patterns of dependency on CDK2^31^ and could benefit from targeted CDK2 inhibition. Expanding this research to other cancer types could elucidate broader applications of CDK2 inhibitor sensitivity, potentially informing new treatment paradigms.

The correlation between high p16 expression and poor overall survival, coupled with the observed sensitivity to CDK2 inhibitors, points to a potential clinical opportunity. Given that p16-high ovarian tumors respond poorly to standard chemotherapy, the lack of adaptation to CDK2 inhibition in p16-expressing cells underscores the potential of CDK2 inhibitors as a targeted therapeutic option. The potential for p16 to become a marker for selecting patients who would benefit from CDK2 inhibitors highlights the importance of mechanism-driven molecular profiling in optimizing cancer treatment strategies.

## Methods

### Cell Culture

Kuramochi and MCF10A cells were seeded at least 48 h prior to imaging in phenol-red-free full-growth medium in glass-bottom 96-well plates (CellVis, P96-1.5H-N) that were coated with collagen prior to seeding. OVCAR3 cells were seeded on µClear-bottom 96-well plates (Greiner, 655090) that were coated with collagen prior to seeding. The seeding density was chosen such that the cells would remain sub-confluent until the end of the imaging period. Cells were first imaged for 24h in full-growth media without drug. The movie was then briefly paused, and the full-growth media was replaced with full-growth media containing drug at the desired concentration. The plate was then re-inserted into the microscope and aligned to its prior position and imaging was continued for an additional 24-48 h. Images were acquired every 20 min on a Nikon Eclipse Ti or Ti2 microscope with a 10X 0.45 NA objective in a humidified, 37°C chamber at 5% CO_2_. Exposure times for all movies for all channels were kept under 500 ms per timepoint to minimize phototoxicity.

### Immunofluorescence

Cells were fixed for 15 min with freshly prepared 4% paraformaldehyde, washed twice with PBS, and incubated with a blocking buffer (3% BSA in PBS) for 1 h at room temperature. Permeabilization was carried out using 0.2% Triton X-100 for 15 min at 4°C. Primary antibody was diluted in blocking buffer and incubated with cells overnight at 4C followed by three washes with 1X PBS. Secondary antibodies conjugated to Alexa Fluor 488, Alexa Fluor 546 or Alexa Fluor 647 were incubated for 1 h followed by three washes with 1X PBS. DNA was stained using Hoechst 33342 dye for 10 min at 1:10000 (Thermo Fisher, H3570). Images were acquired on a Nikon TiE microscope with a 10X 0.45 NA objective. DNA content was determined by taking the integrated intensity of each cell’s Hoechst signal. Cells were delineated as 3-4N DNA content by plotting a histogram of DNA content and drawing a threshold and the end of the 2N peak.

### Long-term treatment experiment

OVCAR3, Kuramochi, and MCF10A cells expressing the nuclear marker H2B-mTurquoise and CDK2 activity sensory DHB-mVenus were seeded in a Poly-D lysine-coated 96-well PhenoPlate (PerkinElmer 6055500) at 3,000 cells per well and treated with the indicated drugs the following day and imaged (day 0). Each condition consisted of 6-12 replicate wells and the entire well was imaged. Treatments were refreshed and cells’ H2B-mTurquoise signal was imaged with a 5X objective on an Opera Phenix (PerkinElmer) on days 0, 1, 2, 4, 7, 10, 14, 18, 21, 25. To quantify the number of cells, nuclear segmentation of the H2B signal was performed using the Harmony software (PerkinElmer).

### Western blotting

Lysates were prepared using diluted 4X LDS sample buffer (ThermoFisher NP007) using equal numbers of cells. Proteins were separated using NuPAGE precast polyacrylamide gels (ThermoFisher NP0301). HRP-conjugated antibodies were used for visualization. Protein concentrations were measured using Bradford (Bio-Rad) method. Primary antibodies were detected with goat secondary antibodies directed against mouse or rabbit IgGs and visualized with ECL Western Blot detection solution (GE Healthcare)

### Multiplexed tumor immunofluorescence staining

The ovarian tissue microarray was obtained from IRB approved protocol (BDR-144221) at Roswell Park Cancer Center entitled “Cell Cycle Plasticity in Human Cancers”. It was initially approved 6/30/2021 and has approval through 6/29/2026. Formalin-fixed paraffin-embedded (FFPE) TMA sections were cut and placed on charged slides. Slides were dried at 65 °C for 2 h. After drying, the slides were placed on the BOND RXm Research Stainer (Leica Biosystems) and deparaffinized with BOND Dewax solution (AR9222, Lecia Biosystems). The multispectral immunofluorescent (mIF) staining process involved serial repetitions of the following for each biomarker: epitope retrieval/stripping with ER1 (citrate buffer pH 6, AR996, Leica Biosystems) or ER2 (Tris-EDTA buffer pH9, AR9640, Leica Biosystems), blocking buffer (AKOYA Biosciences), primary antibody, Opal Polymer HRP secondary antibody (AKOYA Biosciences), Opal Fluorophore (AKOYA Biosciences). All AKOYA reagents used for mIF staining come as a kit (NEL821001KT). Spectral DAPI (AKOYA Biosciences) was applied once slides were removed from the BOND. They were cover slipped using an aqueous method and Diamond antifade mounting medium (Invitrogen ThermoFisher). The biomarker panel comprised of the following antibodies, Cytokeratin (AE1/AE3, Agilent DAKO, 1/300, Opal 480), Cyclin E1 (EP435E, Abcam, 1/400, Opal 520), Cyclin D1 (SP4, Invitrogen, 1/100, Opal 570), RB (4H1, Cell Signaling, 1/150, Opal 620), CDK2 (E8J9T, Cell Signaling, 1/100, Opal 650), Geminin (EPR14637, Abcam, 1/200, Opal 690), p16INK4A (6H12, Leica Biosystems, RTU, Opal 780). Following the staining, slides were imaged on the PhenoImager HT® Automated Quantitative Pathology Imaging System (AKOYA Biosciences). Further analysis of the slides was performed using inForm® Software v2.6.0 (AKOYA Biosciences)

### Tissue recognition and Segmentation

The inForm® Software v2.6.0 was used, which contains feature recognition algorithms to allow automatic identification of specific tissue types based on tissue morphology. Areas of tumor, stroma, and other (no tissue) were marked by extrapolation to segment the entire set of tissues in the tissue microarray. AE1AE3, DAPI, and Autofluorescence were the three channels used to train the segmentation to distinguish tumor from stroma. This enables the Inform software to use nuclear size and shape, cell density, and cytokeratin staining to determine the tumor region. The Autofluorescence channel gives the software clear information about where there is and is not tissue. Cellular segmentation is based on DAPI staining with minimum nuclear size and minimum nuclear intensity.

### Cell tracking and single-cell CDK2 activity analysis

Cell tracking was performed using published MATLAB scripts. Asynchronously cycling cells that divided and received drug treatments during the imaging period were initially grouped into categories based on their time of anaphase relative to the time of drug addition. These cells were then further subcategorized based on their DHB cytoplasmic/nuclear (C/N) ratio after the mitotic event. The C/N ratio was calculated by quantifying the ratio of cytoplasmic to nuclear mean DHB fluorescence. The cytoplasmic component was measured as the mean of the top 50th percentile of a ring of pixels outside of the nuclear mask. Cells were classified as CDK2^inc^ if the DHB C/N ratio was above 0.5 units 3 hours after anaphase; otherwise, they were classified as CDK2^low^. CDK2^inc^ cells are plotted throughout. The median of the single-cell traces within a subcategory was then used to create a median trace with a 95% confidence interval representative of that subcategory.

### Dimensionality Reduction Using UMAP

Uniform Manifold Approximation and Projection (UMAP) was performed to reduce the dimensionality of the dataset using the uwot R package (version 0.2.2)^50^. The input dataset consisted of 435,161 single cells with 168 features extracted from the tissue staining for each cell. Prior to applying UMAP, the data were standardized using the scale() function in R. After scaling, all single-cell features of the data were used as UMAP input parameters. UMAP was implemented using the default parameters provided by the uwot package, which include n_neighbors = 15, min_dist = 0.1, and metric = “euclidean”. The resulting two-dimensional UMAP embeddings were used for clustering and visualization. All analyses were performed in R (version 4.4.1).

### Statistics

For all time-lapse imaging quantifications where the median DHB cytoplasmic/nuclear (C/N) ratio is plotted, 95% confidence intervals are shown as shaded bands for each time point. Non-overlapping shading between treatment groups indicates a statistically significant difference with a p-value < 0.05. Significance levels for unpaired t-tests are reported as p-value summaries on indicated bar charts < 0.05 (*), < 0.01 (**), < 0.001 (***) and < 0.0001 (****) with corresponding star notations. Throughout the manuscript, ‘ns’ denotes no statistical significance. All data shown are representative of multiple experiments. See Supplementary Table 1 for statistics on sample sizes and independent experimental repeats.

## Data Availability

The data that support the findings of this study are available from the corresponding author upon reasonable request. No original code is reported in this paper.

## Acknowledgements

We thank members of the Spencer Lab and Stephen Dann for helpful discussions. B.F. was supported in part by an F31 pre-doctoral fellowship (F31CA284877). E.S.K. and A.K.W. are supported by a grant from NCI (R01 CA247362 and R01 CA267467). This work was funded by an R01 from NIA (1R01AG082942) and an R01 from NIGMS (1R01GM152642) to S.L.S. This work was also supported in part by a Mark Foundation Emerging Leader Award (MFCR-AWD-20-08-0164) and a Damon Runyon Cancer Research Foundation Rachleff Innovation Award (DRR-68-21) to S.L.S. The long-term imaging work was performed at the BioFrontiers Institute’s Advanced Light Microscopy Core (RRID: SCR_018302). The Revvity Opera Phenix is supported by NIH grant 1S10OD025072. We acknowledge Sidney Mahan and Hanna Rosenheck for tissue staining in the Advanced Tissue Imaging Shared Resource at Roswell Park which is supported National Cancer Institute (NCI) grant P30CA016056.

## Author contributions

C.S. and L.W. performed the time-lapse imaging experiments and cell tracking analyses; C.S and L.W. performed and analyzed immunofluorescence experiments; B.F. and C.S performed UMAP analyses; S.L.S. designed the study, with intellectual contributions from C.S. and L.W.. Patient tumor data was provided by a collaboration with A.K.W. and E.S.K., with staining and imaging directed by A.K.W., and data analyses by J.W. and E.S.K.. C.S. and S.L.S. wrote the manuscript. S.L.S. acquired funding and supervised the project.

---

**Fig. S1.**
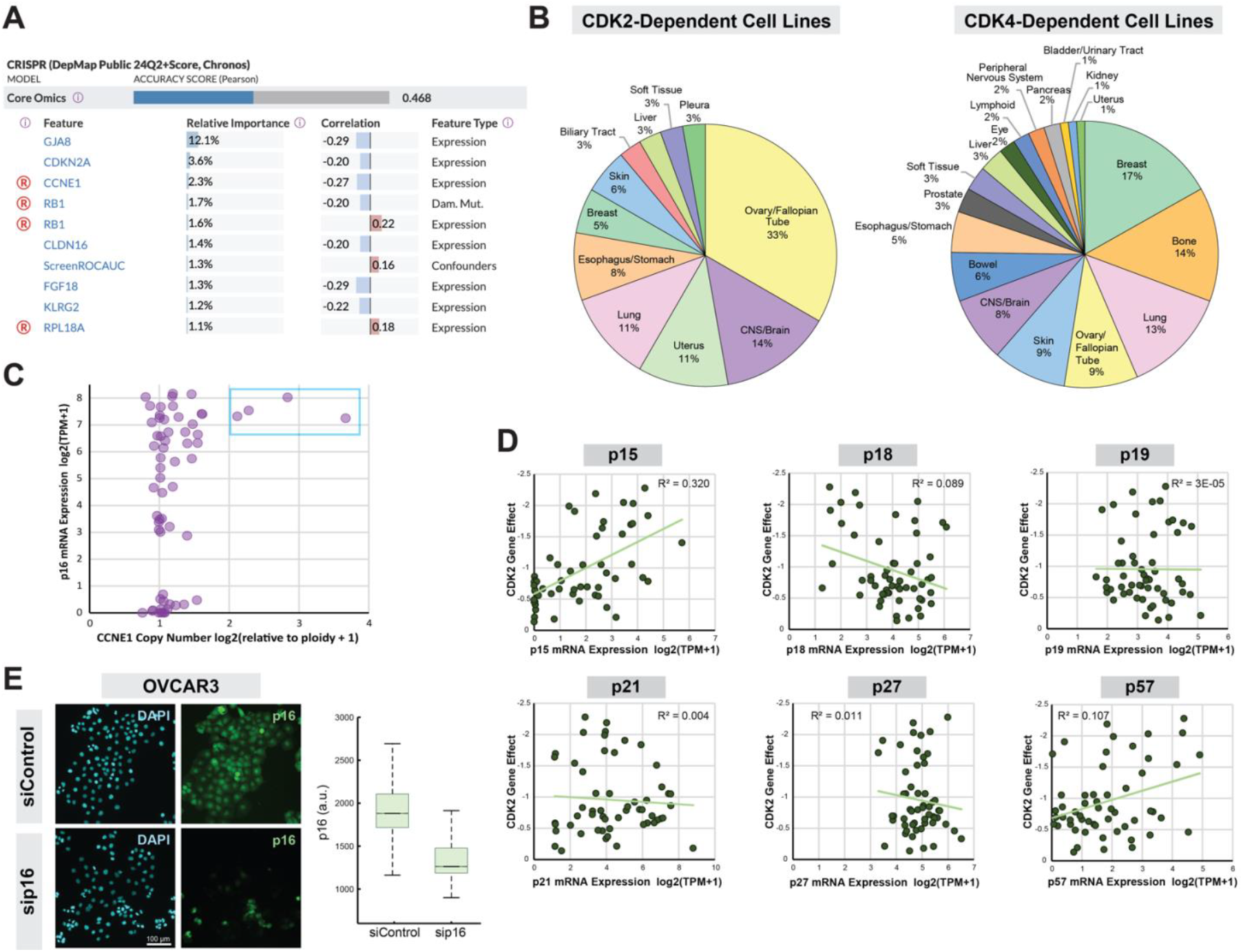
p16 expression is associated with CDK2 dependency in Cyclin E1-amplified ovarian cancers. (**A**) Screenshot from the DepMap predictability tab for CDK2 dependence using CRISPR (DepMap Public 24Q2+Score, Chronos). The relative importance score indicates the impact of a feature on prediction accuracy relative to the other features available to the model (0 to 100% scale) and is calculated using Gini Importance. (Screenshot taken on September 14^th^ 2024). (**B**) Cancer type by CDK dependency (DepMap) defined by having a dependency score less than -1.5 for the indicated gene. (**C**) CCNE1 copy number relative to ploidy by p16 mRNA in ovarian cancers in DepMap. Blue square highlights that all CCNE1-amplified lines have high p16 mRNA. (**D**) DepMap CRISPR Gene Effect score (Chronos) for CDK2 vs mRNA expression for CDKN2B (p15), CDKN2C (p18), CDKN2D (p19), CDKN1A (p21), CDKN1B (p27), and CDKN1C (p57) in ovarian cancers. Linear regression line shown in light green along with R^2^ value. (E) p16 immunofluorescence in OVCAR3 with and without p16 siRNA knockdown for 72h.

**Fig. S2.**
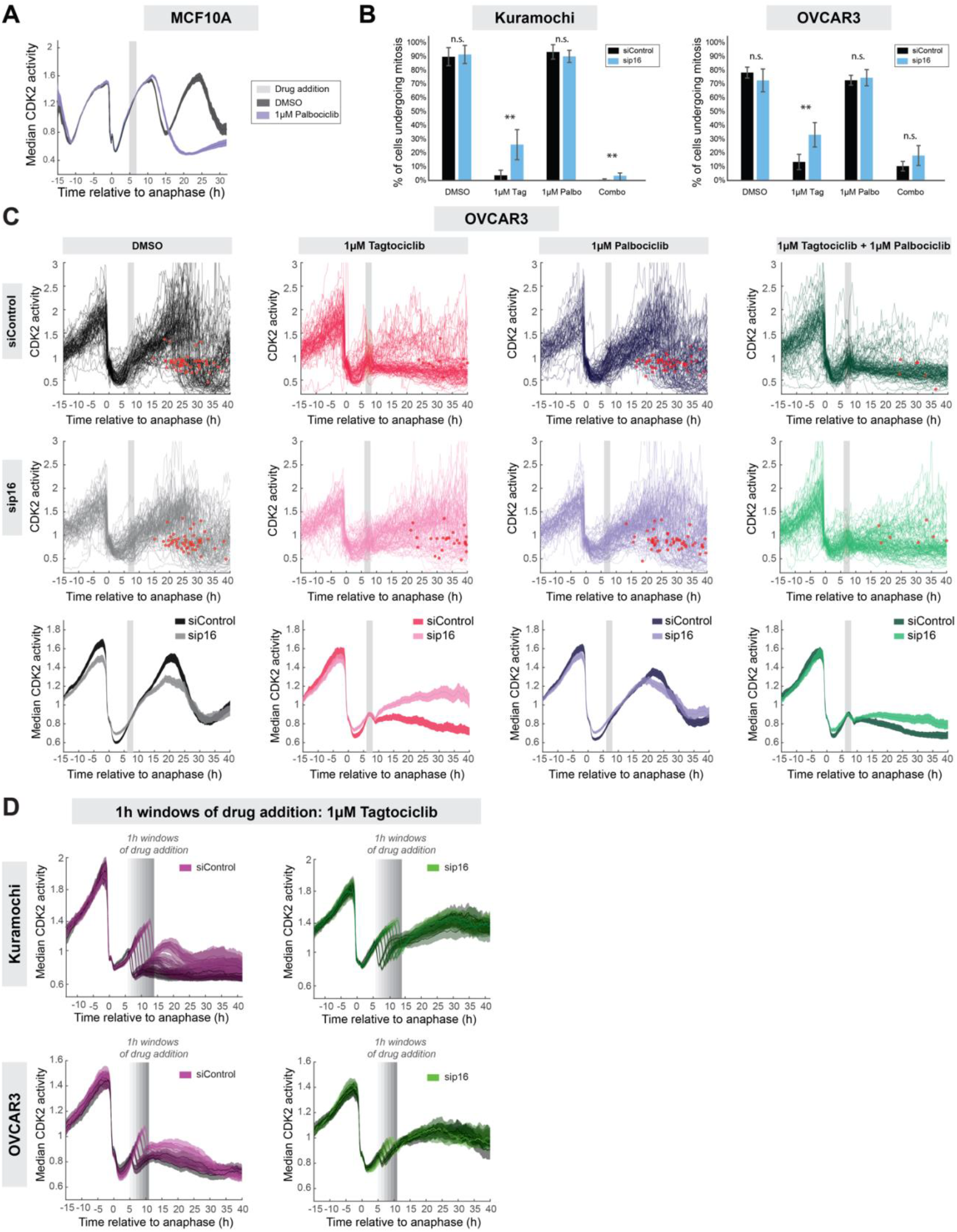
p16 expression modulates the CDK2 activity rebound upon CDK2 inhibition. (**A**) CDK2 activity in MCF10A 2A7T (Cyclin D-mCitrine) cells treated with Palbociclib. Unless stated otherwise, all timelapse plots are median ± 95% confidence interval as shaded bands where non-overlapping shading indicates a statistically significant difference with a p value < 0.05. (**B**) Mitosis count from live-cell movies (red dots) from cells that received drug 5-9 hours after mitosis. Mean of at least 6 technical replicates plotted ± s.d. (**C**) Top two rows: Single-cell traces of CDK2 activity in OVCAR3 cells treated with the indicated drugs. Traces are computationally aligned to the point of anaphase and selected for plotting if they received drug 6-8 h after anaphase (gray bar). Red dots indicate mitosis. Bottom row: 95% confidence interval for the respective plots above. (**D**) 95% confidence interval for OVCAR3 and Kuramochi cells treated with 1μM Tagtociclib at multiple 1h windows of drug addition (gray bars). (Same as Fig. 2E, but with the plots separated for clarity rather than overlaid).

**Fig. S3.**
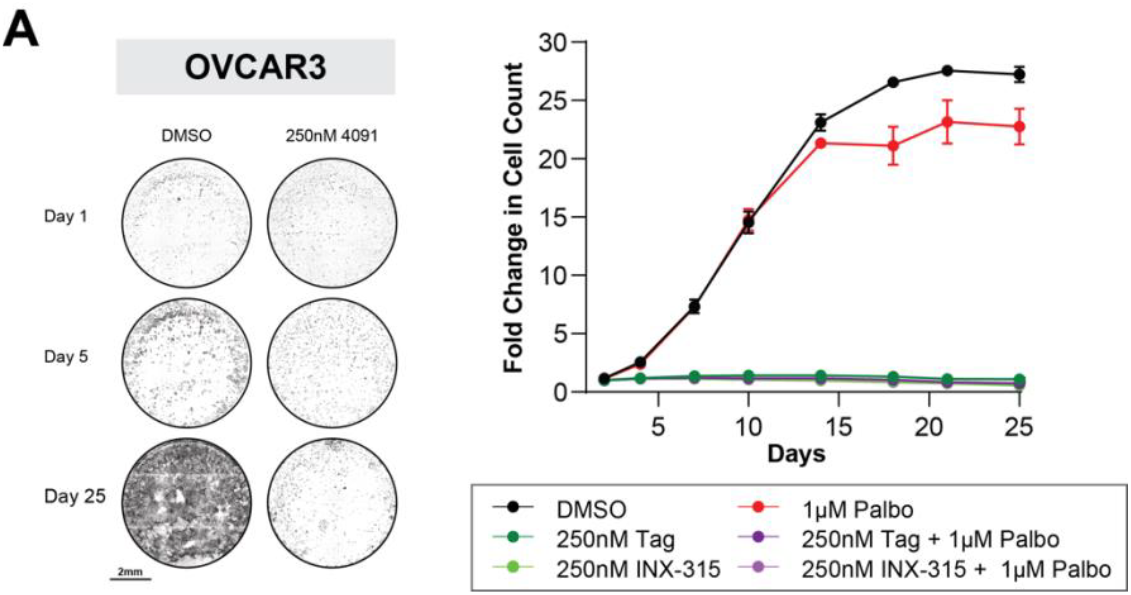
p16 expression modulates CDK2 substrate phosphorylation and cell outgrowth upon CDK2 inhibition. (**A**) Left, long-term treatment of OVCAR3 cells with the indicated drugs. Representative images from Day 0, 5, and 30 for the indicated drug conditions. Right, quantification of images. Mean of at least 3 technical replicates plotted ± s.d.

**Fig. S4.**
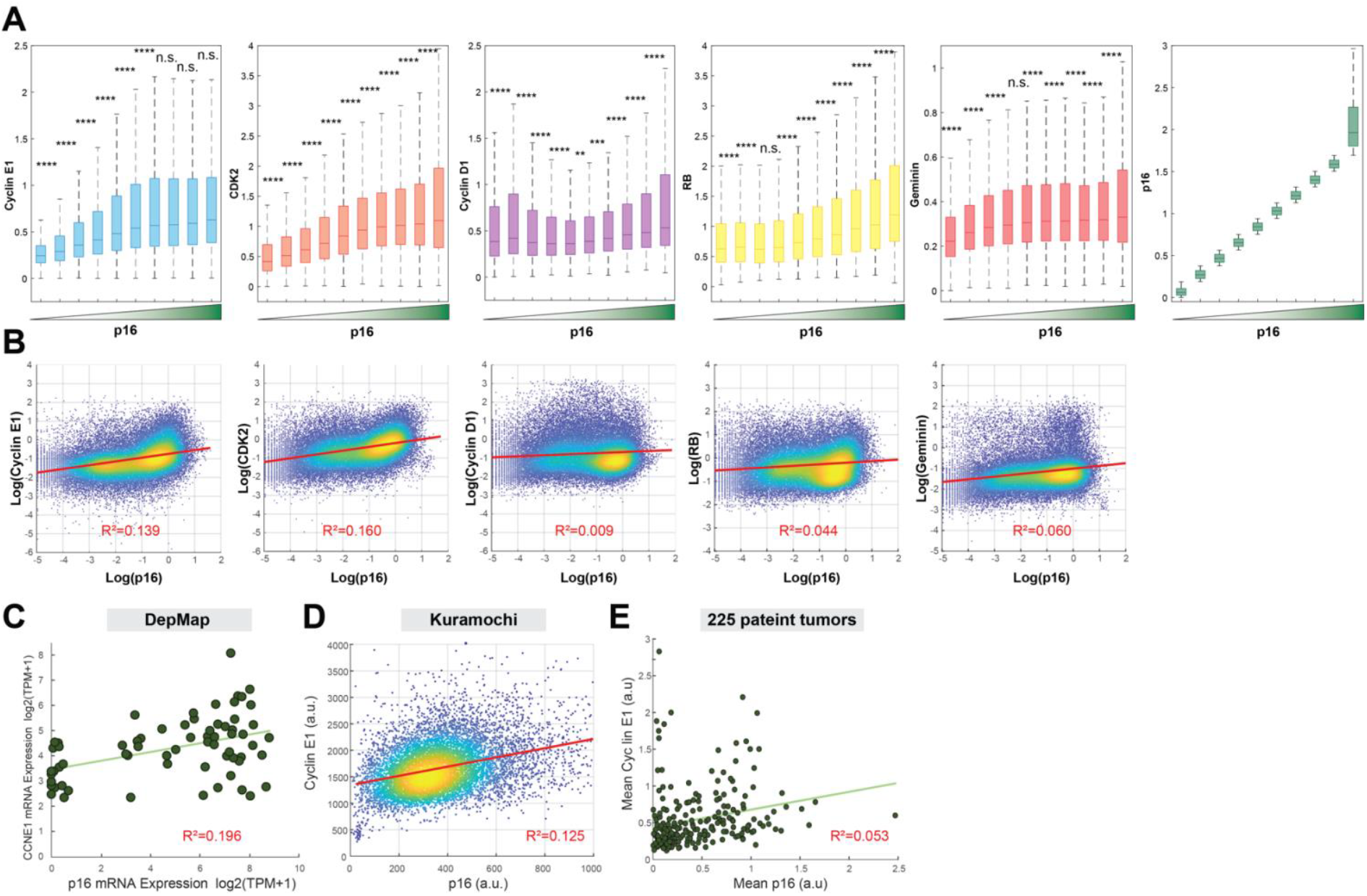
p16 levels are correlated with Cyclin E1 levels. (**A**) Pooled cells from all tumors binned by increasing p16 intensity (right-most plot). The intensity of each marker is plotted for each bin of increasing p16. Error bars represent ± s.d. (**B**) Intensity of each marker in the dataset vs. p16 intensity in all cells in the tumor dataset. The linear regression line is shown in red along with R^2^ values. (**C**) DepMap CCNE1 mRNA expression vs. CDKN2A mRNA expression in ovarian cancers. Linear regression line shown in light green along with R^2^ value. (**D**) Cyclin E1 vs. p16 staining in untreated Kuramochi cells. Linear regression line shown in red along with R^2^ value. (**E**) Mean Cyclin E1 intensity vs. mean p16 intensity for each of 225 tumors. Linear regression line shown in light green along with R^2^ value.

